# Multiplex genomic edition in *Ashbya gossypii* using CRISPR-Cpf1

**DOI:** 10.1101/834218

**Authors:** Alberto Jiménez, Birgit Hoff, José Luis Revuelta

## Abstract

The CRISPR/Cas technologies constitute an essential tool for rapid genome engineering of many organisms, including fungi. The CRISPR/Cas9 system adapted for the industrial fungus *Ashbya gossypii* enables the efficient genome editing for the introduction of deletions, insertions and nucleotide substitutions. However, the Cas9 system is constrained to the existence of an specific 5’-NGG-3’ PAM sequence in the target site.

Here we present a new CRISPR/Cas system for *A. gossypii* that expands the molecular toolbox available for microbial engineering of this fungus. The use of Cpf1 nuclease from *Lachnospiraceae bacterium* allows to employ a T-rich PAM sequence (5’-TTTN-3’) and facilitates the implementation of a multiplexing CRISPR/Cpf1 system adapted for *A. gossypii*. The system has been validated for the introduction of large deletions into five different auxotrophic marker genes (*HIS3, ADE2, TRP1, LEU2* and *URA3*). The use of both crRNA and dDNA arrays in a multi-CRISPR/Cpf1 system was demonstrated to be an efficient strategy for multiplex gene deletion of up to four genes using a single multi-CRISPR/Cpf1 plasmid. Our results also suggest that the selection of the target sequence may significantly affect to the edition efficiency of the system.

## Introduction

*A. gossypii* is a filamentous fungus that is currently exploited in the industry for the production of riboflavin [1]. Furthermore, *A. gossypii* has been proposed as a microbial cell factory for the production of other relevant compounds such as folic acid, nucleosides, recombinant proteins, γ-lactones and biolipids [2–6]. In this context, the development of novel molecular tools is essential for the implementation of rational system metabolic engineering approaches in *A. gossypii*. Hence, a complete toolbox for genomic manipulation is available for *A. gossypii*, including a CRISPR/Cas9 system adapted for this fungus [7–9].

The CRISPR/Cas9 systems have emerged as the foremost technique for genome engineering of many organisms, including yeasts and fungi, with applications that go further beyond the single gene modification (deletions and nucleotide substitutions) [10]. Thus, gene regulation and systems metabolic engineering approaches have been described using CRISPR/Cas9 systems in different yeasts and fungi [10,11].

The CRISPR/Cas9 system of *A. gossypii* shows a high editing efficiency for the introduction of gene deletions, insertions and nucleotide substitutions, which largely facilitates the genomic engineering of the fungus in a marker-less manner [8]. The efficiency of the system in a multinucleated syncytium such as the *A. gossypii* mycelia relies on a one-vector strategy that comprises the expression modules for the CAS9 and the synthetic guide RNA (sgRNA), and the donor DNA (dDNA). The expression of the sgRNA is driven by regulatory sequences from the *A. gossypii SNR52* gene, which is transcribed by RNA Polymerase III.

The CRISPR/Cas-mediated genomic edition have largely transformed the microbial engineering approaches. However, there are some limitations of these technologies regarding the editing efficiency variation between genomic sequences, the off-target effects and the PAM sequence restrictions [12]. Hence, the *A. gossypii* CRISPR/Cas9 system is also restricted to the presence of a 5’-NGG-3’ PAM sequence on the genomic target to generate a double-strand break. Also, multiplexing engineering for the simultaneous edition of different targets and metabolic pathways has not been described for *A. gossypii*.

Cpf1 (recently renamed as Cas12a) is a class 2/type V RNA-guided endonuclease discovered in several bacterial genomes and one archaeal genome [13,14]. Cpf1-mediated genome editing has been described in bacteria, yeasts, plants, insects and vertebrates, including human cells (see [12] and references therein). Cas9 and Cpf1 differ on their evolutionary origin and also show significant structural differences, resulting in different molecular mechanisms and genome editing features [12]:

a. Cpf1 recognizes T-rich PAM sequences, i.e., 5’-TTTN-3’ (AsCpf1, LbCpf1) and 5’-TTN-3’ (FnCpf1), in contrast to the G-rich PAM sequence (5’-NGG-3’) of Cas9 [14].
b. Cpf1-PAM sequences are located at the 5’ end of the target DNA sequence, instead of the 3’ end for Cas9-PAM sequences.
c. Cpf1 cleaves DNA after the +18/+23 position of the PAM, thus creating a staggered DNA overhang, whereas Cas9 cleaves DNA close to its PAM after the −3 position of the protospacer at both strands and creates blunt ends.
d. Cpf1 is guided by a single crRNA and does not require a tracrRNA, resulting in a shorter gRNA sequence than the sgRNA used by Cas9.
e. Cpf1 displays an additional ribonuclease activity that functions in crRNA processing [15]. This might simplify multiplex genome editing, as demonstrated by Zetsche [16] who used a single crRNA array to simultaneously edit up to four genes in mammalian cells.

A G-rich PAM sequence corresponding to the Cas9 can be frequently found in the *A. gossypii* genome, which shows a GC content of 52% [17]. However, the use of CRISPR/Cas9 can be challenging for the genomic edition of AT-rich regions, where the use of Cpf1 nuclease has been shown to be more effective [12]. Here we report a CRIPR/Cpf1 system adapted for the industrial fungus *A. gossypii*. The use of a Cpf1 endonuclease allows implementing a multiplex genome editing system that was efficient to simultaneously edit the auxotrophic markers *HIS3, ADE2, TRP1, LEU2* and *URA3*.

## Materials and Methods

### A. gossypii strains and growth conditions

The *A. gossypii* ATCC 10895 strain was used as a wild-type strain. *A. gossypii* cultures were carried out at 28°C in MA2 rich medium [18]. Auxotrophic mutants were analyzed in SMM complete minimal media lacking the corresponding nutritional requirement (SMM-his, SMM-ade, SMM-trp, SMM-leu, SMM-ura) [19]. *A. gossypii* transformation, sporulation and spore isolation were performed as described previously [18]. Geneticin (G418) (Gibco-BRL) was used where indicated at concentrations of 250 mg/L.

### Assembly of the CRISPR/Cpf1 system for A. gossypii

The CRISPR/Cpf1 system was assembled in a single vector containing all the required modules for genomic editions. The *A. gossyppi* CRISPR/Cas9 vector was used as a backbone that included the replication origins (yeast 2μ and bacterial ColE1) and the resistance markers (Amp^R^ and G418^R^) [8]. The donor DNA and the modules for the expression of Cpf1 and crRNAs were assembled as follows: a synthetic codon-optimized ORF of the Cpf1 enzyme from *Lachnospiraceae bacterium* (LbCpf1) with a SV40 nuclear localization signal was assembled with the promoter and terminator sequences of the *A. gossypii TSA1* and *ENO1* genes, respectively. The expression of the crRNAs was driven by the promoter and terminator sequences of the *A. gossypii SNR52* gene, which is transcribed by RNA Polymerase III. Synthetic donor DNAs comprising the corresponding genomic editions were also assembled in the CRISPR/Cpf1 vector (Fig. 1A). The assembly of the fragments was achieved following a Golden Gate assembly method as previously described [9]. Briefly, a directional cloning strategy was used, by introducing *Bsa*I sites at the ends of the fragments. The *Bsa*I sites are flanked by sequences of 4-nucleotide (nt) sticky ends. Hence, after *Bsa*I digestion, all the modules contain compatible 4-nt sticky ends that facilitate a single-step directional assembly of the CRISPR/Cpf1 vector. The sequences of all the modules are described in Supplementary Material Table S1.

**Figure 1.**
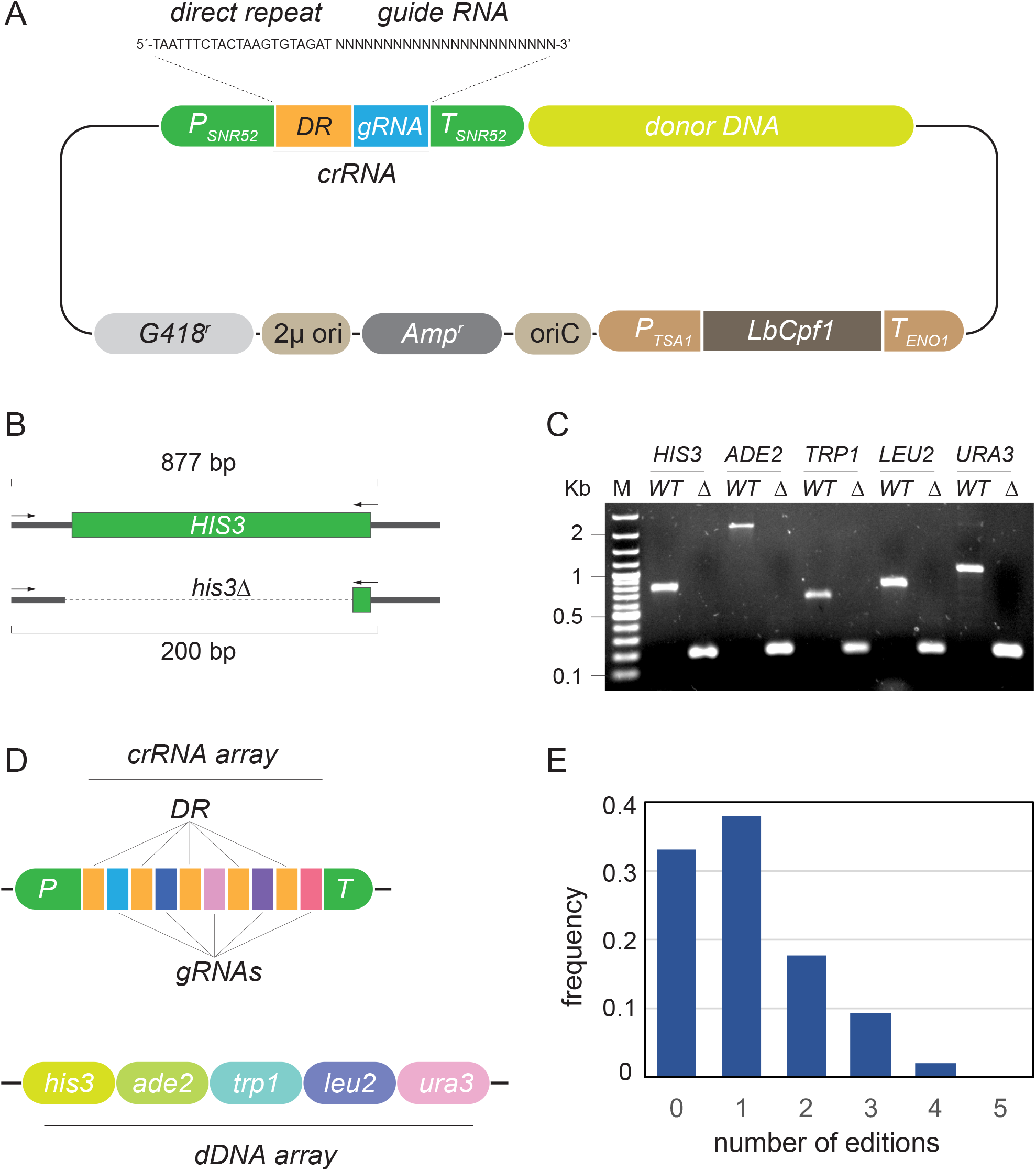
The CRISPR/Cpf1 system adapted for *A. gossypii*. A, modular design of the CRISPR/Cpf1 vector. B, strategy for analytical PCR of homokaryotic clones after CRISPR/Cpf1-HIS3 edition. The PCR primers were designed to amplify different fragments from the WT *HIS3* (877 bp) and the *his3*Δ (200 bp) loci. The same strategy was used for the *ADE2, TRP1, LEU2* and *URA3* loci. C, analytical PCR of homokaryotic auxotrophic clones after the CRISPR/Cpf1 edition. The size of the PCR products correspond to wild-type (WT) and edited (Δ) colonies, respectively. D, organization of the crRNA array and the dDNA array in the multi-CRISPR/Cpf1 vector. Each fragment of the dDNA array is used for HR-directed repair of DSBs generated by Cpf1 in the corresponding loci. E, frequency of genomic editions obtained with the multi-CRISPR/Cpf1 system.

### CRISPR/Cpf1 genome editing in A. gossypii

Spores of indicated strains of *A. gossypii* were transformed with 5-15 μg of the corresponding CRISPR/Cpf1 plasmid. Heterokaryotic transformants were selected in G418-containing MA2 media. The G418 resistant colonies were isolated and grown in G418-MA2 media during 2 days to facilitate the occurrence of genomic edition events. The loss of the CRISPR/Cpf1 plasmids, which is essential to avoid the genomic integration of the plasmid, was carried out after the sporulation of the heterokaryotic clones in SPA media lacking G418. Homokaryotic clones were isolated in MA2 media lacking G418. The genomic editions leading to auxotrophic colonies were analysed both by auxotrophic screening and analytical PCR. All the genomic editions were further confirmed by DNA sequencing of the analytical PCR fragments.

## Results and discussion

### Design of a CRISPR/Cpf1 system adapted for *A. gossypii*

The CRISPR/Cas9 tool for *A. gossypii*, which was designed as a one-vector system, is optimized for the genomic engineering of multinucleated germlings of the fungus [8]. In order to expand the repertoire of genomic editing tools for *A. gossypii*, a CRISPR/Cpf1 system has been designed. As mentioned above, Cpf1 has several differences with Cas9 and, therefore, the development of a CRISPR/Cpf1 system adapted for *A. gossypii* provides an additional mechanism for the genomic edition of this industrial fungus.

The Cpf1 enzyme from *Lachnospiraceae bacterium*, which recognizes the PAM sequence 5’-TTTN-3’, was used and its expression was driven by the promoter and terminator sequences of the *A. gossypii* genes *TSA1* and *ENO1*, respectively. The Cpf1 module was assembled with functional sequences of the CRISPR/Cas9 vector that was used as a backbone [8] (Fig. 1A). The crRNAs comprised a 21-bp direct repeat (DR) sequence together with a target-specific 23-bp sequence that works as the guide RNA (gRNA) for the ribonucleoprotein Cpf1/crRNA (Fig. 1A). In addition, the CRISPR/Cpf1 system comprised a sequence that functions as donor DNA (dDNA) for the repair of DNA double strand breaks (Fig. 1A).

### CRISPR/Cpf1-mediated single gene editing of *A. gossypii*

The functionality of the CRISPR/Cpf1 system was assessed for the ability to generate large gene deletions that cause auxotrophic phenotypes. Thus, the screening of genomic editions was carried out by analyzing the corresponding nutritional requirements in selective culture media. Five different genomic targets were selected for single CRISPR/Cpf1-mediated editing events: *ADL270C* (*HIS3*), *ACR210C* (*ADE2*), *AER014W* (*TRP1*), *AAL012C* (*LEU2*) and *AEL059W* (*URA3*).

Hence, synthetic DNA sequences for the corresponding crRNAs and dDNAs modules were designed and assembled by Golden Gate cloning in the CRISPR/Cpf1 vector (sequences are described in Supplementary Material Table S1). The five CRISPR/Cpf1 plasmids were used to transform spores of the wild-type strain of *A. gossypii* and positive heterokaryotic clones were selected in G418-containing medium. 250 homokaryotic clones were isolated after sporulation of the primary heterokaryotic clones and the edition efficiency of each plasmid was calculated by auxotrophic screening in selective media (Table 1). The five CRISPR/Cpf1 plasmids were able to produce the corresponding edition event, which was confirmed by analytical PCR (Fig. 1B-C) and DNA sequencing of the amplicons (not shown). However, the editing efficiency of the CRISPR/Cpf1 plasmids differed substantially depending of the genomic target sequence (Table 1). While both *HIS3* and *ADE2* editions exhibited high editing efficiencies (68.4% and 77.2%, respectively), the plasmid for CRISPR/Cpf1-*TRP1* deletion showed a 19.2% of editing efficiency. This variation can be explained by differences either in the recognition of the PAM sequence or in the specificity of the target DNA binding and cleavage of each crRNA, as previously described for other CRISPR/Cas9/Cpf1 systems [12]. Indeed, significant variability has already been reported in the CRISPR/Cas9 editing efficiency of the *A. gossypii ADE2* gene depending of the sequence of the sgRNA used [8]. Also, the low editing efficiency shown at the *TRP1* locus could be related to the presence of the complex chromatin structure of a neighbor centromere.

**Table 1.** Editing efficiency of the CRISP-LbCpf1 systems.

| Edition event | Editing efficiency (%)<sup>1</sup> |  |  |  |
| --- | --- | --- | --- | --- |
|  | LbCfp1 | LbCfp1-crRNA | LbCfp1-crRNA-dDNA | LbCfp1-crRNA array-dDNA array |
| his3<sup>-</sup> | - | - | 68.4 | 19.2 |
| ade2<sup>-</sup> | - | - | 77.2 | 29.6 |
| trp1<sup>-</sup> | - | - | 19.2 | 10.4 |
| leu2<sup>-</sup> | - | - | 62.2 | 24 |
| ura3<sup>-</sup> | - | - | 58.8 | 25.6 |

<sup>1</sup> Editing efficiency was calculated by auxotrophic screening of 250 homokaryotic colonies isolated after sporulation of primary heterokaryotic transformants.

### CRISPR/Cpf1 multiplex genome editing using a single crRNA array

Multiplex genome editing can be achieved with CRISPR/Cas9 systems, but it requires more complex sgRNA modules than those of the CRISPR/Cpf1 systems, due to the intrinsic Cpf1 ribonuclease activity that facilitates crRNA processing [16]. Hence, a crRNA array was designed for the multiplex gene deletion of the five auxotrophic markers *HIS3, ADE2, TRP1, LEU2* and *URA3* using a multi-CRISPR/Cpf1 vector. The crRNA array comprised specific gRNAs for each target sequence preceded by the direct repeats that enable the formation of five different crRNAs to drive the Cpf1 nuclease activity (Fig. 1D). In addition, the multi-CRISPR/Cpf1 plasmid contained an array of dDNA sequences for HR-directed repair of the double-strand breaks generated by Cpf1 in the target genes (Fig. 1D). In contrast to the singleplex CRISPR/Cpf1 plasmid, the dDNA array was assembled between the oriC and the Cpf1-expression module in the multi-CRISPR/Cpf1 plasmid.

The multi-CRISPR/Cpf1 plasmid was used for transformation of spores of the wild-type strain of *A. gossypii*. Primary G418-resistant heterokaryotic colonies were isolated in selective medium, thereby confirming the functionality of the plasmid. Then, the isolation of homokaryotic clones was carried out after the sporulation of heterokaryotic transformants. The frequency of genomic editions was evaluated in 250 homokaryotic clones by auxotrophic screening in selective media (Table 2). The presence of the genomic editions was further confirmed by analytical PCR and DNA sequencing (not shown). Our results revealed a significant reduction in the editing efficiency of each genomic sequence (Table 1). As a consequence, one third of the homokaryotic clones analyzed were devoid of auxotrophies. However, the multi-CRISPR/Cpf1 demonstrated a satisfactory efficiency for the recovery of double and triple mutants (44 and 23 out of 250, respectively) (Fig. 1D). Although the multi-CRISPR/Cpf1 system was challenged with the generation of up to five genomic editions, quintuple auxotrophic mutants could not be detected among the analyzed colonies. However, albeit at low frequency, quadruple mutants were isolated, thus demonstrating that the multi-CRISPR/Cpf1 system can be applied for multiplex genomic edition approaches (Fig. 1E and Table 2).

**Table 2.**
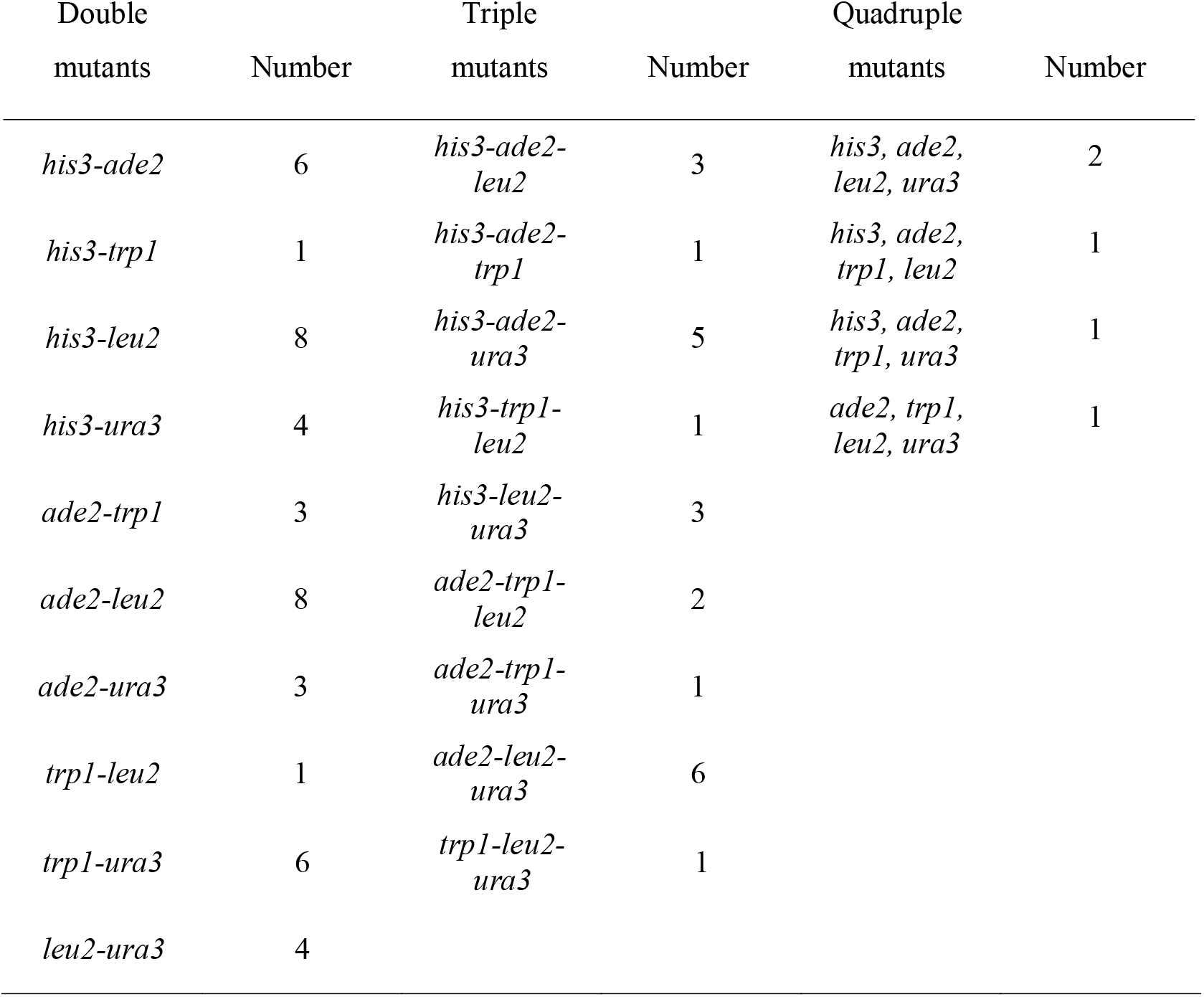
Recovery of multiple mutants with the multiplex CRISP-LbCpf1 system.

| Double mutants | Number | Triple mutants | Number | Quadruple mutants | Number |
| --- | --- | --- | --- | --- | --- |
| *his3-ade2* | 6 | *his3-ade2-leu2* | 3 | *his3, ade2, leu2, ura3* | 2 |
| *his3-trp1* | 1 | *his3-ade2-trp1* | 1 | *his3, ade2, trp1, leu2* | 1 |
| *his3-leu2* | 8 | *his3-ade2-ura3* | 5 | *his3, ade2, trp1, ura3* | 1 |
| *his3-ura3* | 4 | *his3-trp1-leu2* | 1 | *ade2, trp1, leu2, ura3* | 1 |
| *ade2-trp1* | 3 | *his3-leu2-ura3* | 3 |  |  |
| *ade2-leu2* | 8 | *ade2-trp1-leu2* | 2 |  |  |
| *ade2-ura3* | 3 | *ade2-trp1-ura3* | 1 |  |  |
| *trp1-leu2* | 1 | *ade2-leu2-ura3* | 6 |  |  |

|  |  |  |  |
| --- | --- | --- | --- |
| *trp1-ura3* | 6 | *trp1-leu2-*<br>*ura3* | 1 |
| *leu2-ura3* | 4 |  |  |

*A. gossypii* lacks a known sexual cycle and, therefore, the construction of strains combining different gene modifications by sexual crossing is not possible. Hence, the construction of engineered strains containing multiple gene modifications relies on successive cycles of marker-mediated DNA integration followed by marker removal [20]. Although effective, this is a very labor and time-consuming procedure. Multiplex gene modification methods, as demonstrated in this work, enables the modification of several genes simultaneously and permits a rapid generation of engineered strains.

CRISPR/Cas9 technologies has been described for genome editing of many filamentous fungi using different strategies for vector construction and transformation methods [11]. In contrast, Cpf1-based systems have been only described for two Aspergilli species among filamentous fungi [21]. The Cpf1-based genomic edition of Aspergilli required the co-transformation of the Cpf1-expressing vector with an oligonucleotide (dDNA) for genomic-site-directed mutagenesis [21]. The CRISPR/Cpf1 system presented in this work is fully assembled as a one-vector system containing all the modules (Cpf1, gRNA and dDNA), which facilitates the transformation events in multinucleated cells, as previously described for the CRISPR/Cas9 system [8]. In addition, the multi-CRISPR/Cpf1 system represents the first Cpf1-based system for multiplex genomic edition so far. Hence, the use of genome-scale gRNA libraries may constitute a powerful tool for the discovery and functional annotation of genetic elements that modulate transcriptional activity in *A. gossypii*. In this regard, chromatin accessibility as well as target sequence composition largely affects Cpf1 activities [22]. Thus, the development of Cpf1 activity prediction algorithms considering these two factors would substantially improve the selection of target sequences and the efficiency of multi-CRISPR/Cpf1systems.

## Conclusions

A versatile CRISPR/Cpf1 system has been adapted for the efficient genome editing of *A. gossypii*. This technology complements the current CRISPR/Cas9 system, thus minimizing the limitations associated to the existence of specific PAM sequences within the target sites. In addition, a multi-CRISPR/Cpf1 has been designed and validated for multiplex gene deletion of four auxotrophic markers. Hence, the existence of two CRISPR/Cas systems adapted for microbial engineering of *A. gossypii* will largely contribute to facilitate system metabolic engineering approaches in this industrial fungus.

## Acknowledgements

This work was financed by grants from the Spanish Ministerio de Economía y Competitividad (BIO2017-88435-R) and Junta de Castilla y León (SA016P17) to JLR and AJ. We thank María Dolores Sánchez and Silvia Domínguez for excellent technical help.

## Conflict of interest

The authors declare no financial or commercial conflict of interest

